# FSI (Fluctuating Selection among Individuals) Reduces the Mean Fixation Time (Generations) of a Mutation

**DOI:** 10.64898/2026.01.21.700920

**Authors:** Xun Gu

**Affiliations:** Department of Genetics, Development and Cell Biology, Iowa State University, Ames, IA 50011, USA; The Laurence H. Baker Center in Bioinformatics on Biological Statistics, Iowa State University, Ames, IA 50011, USA; Program of Ecological and Evolutionary Biology, Iowa State University, Ames, IA 50011, USA

## Abstract

A common assumption in molecular evolution is the fixed selection nature of a mutation, for instance, a neutral mutation is selectively neutral for all individuals who carry the mutation, and so forth a deleterious or beneficial mutation. Our recent work challenged this presumption, postulating that individuals with a specific mutation exhibit a fluctuation in fitness, short for FSI (fluctuating selection among individuals). Moreover, an intriguing phenomenon called *selection-duality* emerges, that is, a slightly beneficial mutation could be a negative selection (the substitution rate less than the mutation rate). It appears that selection-duality is bounded: the low-bound is the *generic neutrality* where the mutation is neutral by the means of fitness on average, while the up-bound is the *substitution neutrality* where the substitution rate equals to the mutation rate. In this paper, we conducted a thorough theoretical analysis to evaluate how many generations needed for a selection-duality mutation to be fixed in a finite population. A striking finding is that the mean fixation time of a selection-duality mutant, including the generic neutrality and the substitution neutrality, is approximately identical, which is considerably shorter than the case of strict neutrality without FSI. One may further envisage that the fast-fixation nature of selection-duality mutations could result in a considerable genetic reduction at linked sites.

## Introduction

de Jong et al. (2024) pointed out the reason why the neutralist-selectionist debate (Kimura 1968; 1983) has not been resolved for over half century (Hahn 2008; Hughes 2008; Nei et al. 2010; Kern and Hahn 2018; Jensen et al. 2019; Munoz-Gomez et al. 2021; Gu 2021; Galtier 2024; McCandlish and Stoltzfus 2014): many of the tested predictions could be consistent with both viewpoints with some additional assumptions. Thus, both sides agreed that the existed models need to be examined carefully and the generalization is high desirable. A common assumption in molecular evolution is the fixed selection nature of a mutation, postulating that any genotype (heterozygote or homozygote) of a mutation has the same fitness effect among individuals with the same genotype. That said, a neutral mutation is selectively neutral for all individuals who carry the mutation, and so forth a deleterious or beneficial mutation. Our recent work (Gu 2021; 2025a; 2025b) challenged this presumption: a new population genetics model called FSI, short for the fluctuating selection among individuals, was proposed. In other words, FSI refers to dealing with the phenomenon when individuals with a specific mutation exhibit a broader phenotype variation, resulting in a fitness fluctuation (Gu 2021). Meanwhile, the fitness effect of the wildtype remains a constant.

The biological basis of FSI of mutations is complex. For instance, human geneticists have well-demonstrated that mutations frequently exhibit different effects on individuals, challenging the fixed view of mutational effects (Riordan and Nadeau 2017; Eldar et al. 2009; Raj et al. 2010; Jensen et al. 2025). It can be roughly classified into several categories, such as *genetic background* (Chandler et al. 2013; Mullis et al 2018), *stochastic gene expression* (Raj and van Oudenaarden 2007; Elowitz et al. 2002; Ozbudak et al. 2002; Maamar et al. 2007; Vu et al. 2015), *incomplete penetrance* (Khoury 1988; Eldar et al. 2009; Suel et al. 2007), as well as *the complexity of genotypephenotype map* (Dowell et al. 2010; Lehner 2013; Taylor and Ehrenreich 2014). It should be noted that those categories are not mutually excluded (Raj et al. 2010).

The pattern of molecular evolution and population genetics of FSI has been studied recently. Gu (2025a; 2025b) showed that, intriguingly, a novel phenomenon called ‘selection duality’ emerges from FSI: mutations that are statistically slightly beneficial are subject to a negative selection, which would merge to the conventional strict neutrality when FSI vanishes. Gu (2025a) showed that the substitution rate tends to inversely related to the log of effective population size (*N*_*e*_) when FSI is nontrivial, as observed by Galtier et al. (2016). It appears that the nearly-neutral theory (Ohta 1973; 1993) may be a special of no FSI. Moreover, Gu (2025a) developed a statistical procedure to predict the relative strength of FSI to the *N*_*e*_-genetic drift (Lynch M et al. 2011). Meanwhile, Gu (2025b) studied the population genetics of FSI, and in particular evaluated the effects of FSI on sequence divergence between species and genetic diversity within a population (Sawyer and Hartl 1992), revealing a provocative interpretation for the McDonald-Kreitman test that differs from the neutralist-view or the selectionist-view. It should be noticed that FSI does not exclude the possibility of local adaptation through phenotype plasticity (Sommer 2020; Lee et al. 2022).

In this article, we address a fundamental problem in population genetics: how many generations, on average, is required for a selection-duality mutation to be fixed in a population, as short for the mean fixation time problem. One may see the mathematical introduction from Crow and Kimura (1970) or Ewens (2004). It is well known that the mean fixation time (*T*_*fix*_) of a strict neutral (no FSI) is 4*N*_*e*_ generations, the focus of study turns out to whether *T*_*fix*_ under selection-duality is less than 4*N*_*e*_, or not. Our study may have two important implications. First, it helps to quantify the basic scenario of molecular evolution (Kimura 1983) that within-population diversity is the transition phase of divergence between species. Second, it helps to understand the diversity-reduction effect of selective sweeps at linked nucleotide sites. While selection sweeps have been regarded as signals of adaptive evolution, the empirical evidence has been lacking. The preliminary analysis of Gu (2025a) suggested that the mean fixation time could be much less 4*N*_*e*_, a similar pattern to the positive selection. This raises an interesting hypothesis that fixation of a selection-duality mutation could cause the reduction of genetic diversity at linked sites, without invoking the adaptive notion. We will provide a detailed discussion about this issue.

## Results

### The Wright-Fisher diffusion model under FSI

Consider a random mating population of a monoecioys diploid organism. In a finite population, each individual produces a large number of offspring and that exactly *N* of those survive to maturity. Let *a* and *A* be the mutant and wild-type alleles at a particular locus, respectively, whose fitness effects are additive. The FSI model postulates that the fitness effect of mutant *a* is fluctuating among individuals, whereas that of wild-type *A* remains a constant. Therefore, the relative fitness of genotype *AA, Aa*, or *aa* is, on average, given by 1, 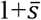 and 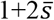, respectively; 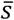 is called the mean of the selection coefficient (*s*) of mutant *a*. Let 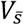 be the variance of 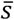 and *Var*(*s*) be the variance of *s*, respectively. Noting that the number of mutant *a* is 2*Nx*, where *x* is the frequency of mutant *a* in a generation, we obtain

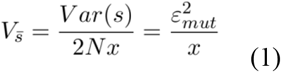

where 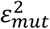 is called the FSI-coefficient; a large value means a strong FSI and *vice versa*; the subscript indicates that FSI is mutation-induced. It should be noticed that the population genetics of fluctuating selection between generations (FSG) has been widely studied (Karlin and Levikson 1974; Karlin and Lieberman 1974); one may also see Gu (2024) for a commentary about its theoretical inconsistency. Eq.(1) makes a distinction of FSI from FSG, as the variation of FSI is inversely dependent of the mutation frequency.

Gu (2025b) developed a Wright-Fisher diffusion model under FSI, where the infinitesimal mean *μ*(*x*) and the infinitesimal variance *σ*^*2*^(*x*) are, respectively, given by

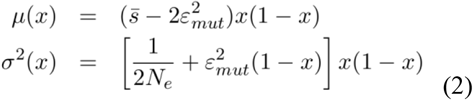

where *N*_*e*_ is the effective population size. Briefly speaking, *μ*(*x*) describes those determinative factors that may influence the gene frequency change, and *σ*^*2*^(*x*) describes the random effect of genetic drifts. In following-up analysis we define the (adjusted) selection-FSI ratio (*ρ*) by

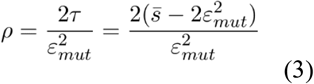

where 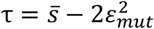. Apparently, *ρ*>*0* guarantees *μ*(*x*)>0, indicating a positive selection, or *ρ<0* indicating a negative selection. Meanwhile, FSI emerges as a new resource of genetic drift, measured by the FSI-strength 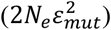. Reading the FSI-strength by 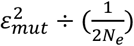, one may claim a dominant FSI-genetic drift when 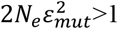, or a dominant *N*_*e*_-genetic drift when 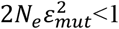. Sometimes it is concise to use a relative measure of FSI-strength (*F*), that is,

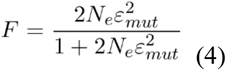

Thus, *F* increase from 0 at 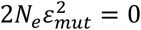 (no-FSI), and approaches to 1 when 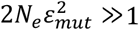.

### Substitution rate and selection duality

Let *v* be the mutation rate and *N* be the census population size. From the view of population, the substitution rate (*λ*) can be defined by the amount of new mutations per generation (2*Nv*) multiplied by the fixation probability of a single mutation with the initial frequency of 1/(2*N*), based on the assumption of rare, single *de novo* mutation event. Let *u*(*p*) be the fixation probability of a mutation in a finite population, with the initial frequency *p*. Formally, the substitution rate can be written by 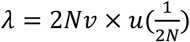.

After some calculations in Methods, we obtain

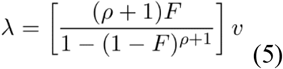

It should be noticed that *λ*>*v* (the substitution rate greater than the mutation rate) when *ρ*>0, indicating a positive selection, whereas *λ*<*v* (the substitution rate less than the mutation rate) when *ρ*<0, indicating a negative selection. In the case of no-FSI, Eq.(5) is reduced to the well-known formula first reported by Kimura (1962).

Further analysis of Eq.(5) indicates that, when 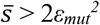, i.e., the mutation is highly beneficial, molecular evolution is mainly driven by a positive selection, and FSI only plays a marginal role. In the same manner, when 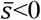, a strong purifying selection becomes dominant. Nevertheless, between those two cases an intriguing phenomenon called *selection-duality* emerges: a slightly beneficial mutation defined by

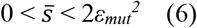

is subject to a negative selection (*λ*<*v* because of *ρ*<0). It appears that selection-duality defined by Eq.(6) is bounded by two types of neutrality. The low-bound is the *generic neutrality* 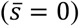, where the mutation is neutral by the means of fitness. On the other hand, the up-bound of selection-duality is the *substitution neutrality* 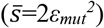, where the substitution rate equals to the mutation rate (*λ*=*v*). The broadness of selection duality depends on the magnitude of 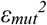. Without FSI, i.e., 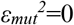, the selection duality vanishes as the generic neutrality and the substitution neutrality merge onto the classical neutrality. In addition to those boundary neutralities, the middle-point of selection-duality at 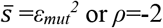, called FSI-neutrality, may play a pivotal role in the new theory of molecular evolution, as shown later.

### Mean fixation time of a new mutation

Let us consider, under FSI, how many generations it will take for a mutant to be fixed in a finite population. To this end, one may trace a particular mutant and study the average number of generations at which the frequency of the allele become 1. Let *T*_*fix*_(*p*) be the mean fixation time of a mutation, given the initial frequency *p*. Theoretically, this average fixation time can be obtained by integrating the sojourn time that the gene frequency spends at a particular value *x*, given that the allele is going to be fixed (Kimura and Ohta 1969). Given Eq.(2), it has been shown that *T*_*fix*_(*p*) can be computed by

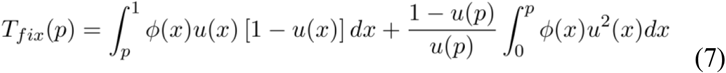

where several variables in Eq.(7) are given by

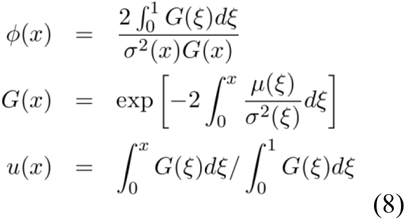

respectively; one may see Methods as well as the Appendix for mathematical introduction.

We are particularly interested in the mean fixation time of a new mutation. For a new mutation arising in the population, one may set the initial frequency *p*=1/(2*N*), where *N* is the census population size. Except for a very small population size, the mean fixation time for a nearly arising mutation can be well approximated by letting *p* → 0. One may write 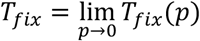, resulting in

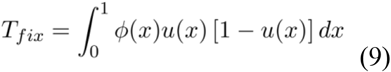

(see the Appendix).

### Mean fixation time in selection duality

Our major goal is to compute the mean fixation time of selection-duality mutation, that is, −4 ≤ *ρ* ≤ 0. In spite that *T*_*fix*_ is not analytical in a general case, we choose three special cases whose analytical forms can be derived, which are sufficient to demonstrate the general pattern. They are the substitution neutrality (*ρ* = 0) as up-boundary, the generic neutrality (*ρ* = −4) as down-boundary, and the FSI-neutrality (*ρ* = −2) as the middle-point of selection duality. Based on Eq.(8), *ϕ*(*x*) are given by

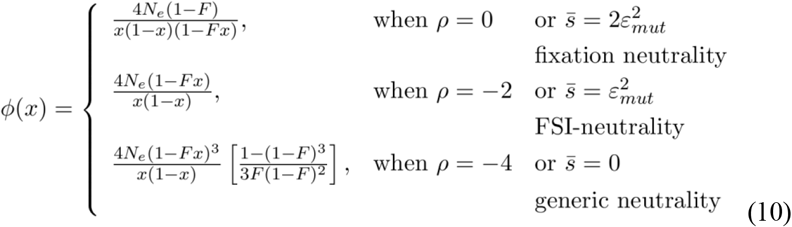

under *ρ* = 0, −2, −4, respectively, as well as *u*(*x*) by

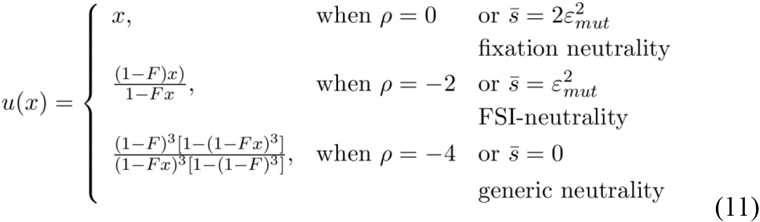

After plugging Eq.(10) and Eq.(11) onto Eq.(9) following by appropriate mathematical treatments, we obtain the following main results.

First, we found that the mean fixation time (*T*_*fix*_) of a new-arising mutation with substitution neutrality (*ρ* = 0) is identical to that with FSI-neutrality (*ρ* = −2), as given by

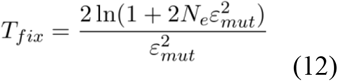

One can easily verify that *T*_*fix*_=4*N*_*e*_ only when 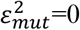; otherwise *T*_*fix*_<4*N*_*e*_ always. Roughly speaking, when 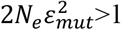, the magnitude of *T*_*fix*_ is determined by the inverse of FSI-coefficient, and increases, in a log-scale, with the increase of effective population size. For any *ρ* between 0 and −2, the analytical form of *T*_*fix*_ is not available, yet the numerical calculation shows that the difference from Eq.(12) is rather marginal.

Second, the mean fixation time (*T*_*fix*_) of a new-arising mutation with generic neutrality (*ρ* = −4) is analytical yet complicated. After some algebras, we obtain

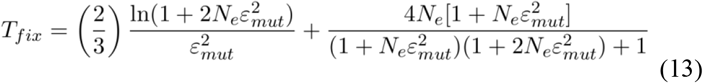

Similar to Eq.(12), we have *T*_*fix*_=4*N*_*e*_ only when 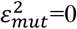; otherwise *T*_*fix*_<4*N*_*e*_ always.

When 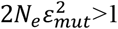, Eq.(13) can be approximated simplified by

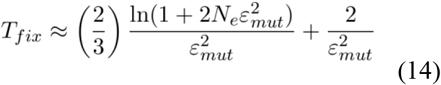

Eq.(14) reveals that the mean fixation time of a newly-arising mutation with generic neutrality has similar patterns to those with substitution neutrality or FSI-neutrality, Indeed, the magnitude of *T*_*fix*_ is determined by the inverse of 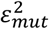. We noticed that, while *T*_*fix*_ increases with *N*_*e*_ in a log-scale, the approaching rate is about one-third of that in the fixation or FSI-neutrality.

### Approximations of *T*_*fix*_ under selection-duality

We have shown that the mean fixation time is identical between substitution neutrality (*ρ*=0) and FSI-neutrality (*ρ*=−2). An interesting question is about the situation when a mutation is under positive selection-duality, i.e., −2< *ρ*<0. Because of the mathematical complexity, the mean fixation time for positive selection-duality cannot be analytically derived. Nevertheless, numerical analysis has shown that the mean fixation time in the case −2< *ρ*<0 is very close. We thus formulated an intuitive approximation of *T*_*fix*_ in the range of −2< *ρ*<0 that satisfies

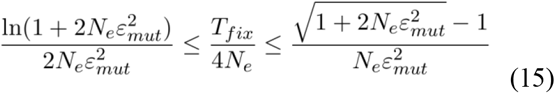

(see the Appendix).

We also notice that the mean fixation time at FSI-neutrality (*ρ*=−2) differs from that at generic neutrality (*ρ*=−4). Though the difference is marginal, it is interesting to evaluate *T*_*fix*_ for any mutation in range of negative selection-duality, i.e., −4< *ρ*<-2. Similar to above, the mean fixation time in negative selection-duality is generally not analytical except for the case of *ρ*=−3. As shown by the Appendix, we proposed an approximate formula for the mean fixation time of a mutation in the range of −4< *ρ*<-2 as follows

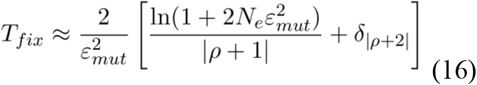

where *δ*_|*ρ*+2|_ is an index function: *δ*_|*ρ*+2|_ =0 when |ρ + 2| = 0, otherwise *δ*_|*ρ*+2|_ =1. It should be noticed, as long as 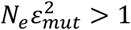, Eq.(16) represents almost the accurate form for *ρ*=−2, −3 or −4, respectively.

### Mean fixation time of a mutation given initial frequency

The mean fixation time of a selection-duality mutation given an initial frequency can be derived in a similar way, though the result may be tedious in algebra. We first claim that *T*_*fix*_(*p*) is identical between the substitution neutrality and the FSI-neutrality. Here we use the substitution neutrality (*ρ* = 0) for illustration. In this case one can show *G*(*x*)=1, *u*(*x*)=*x*, and *ϕ*(*x*) = 2/*σ*^2^(*x*) and therefore,

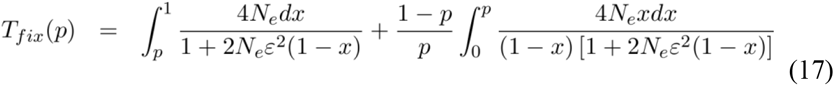

After some calculations, we obtain

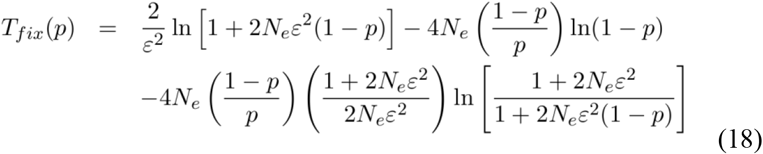

In the case of no FSI, i.e., 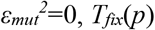, *T*_*fix*_(*p*) is reduced to the result of Kimura and Ohta (1969), that is,

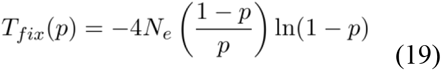

### Mean loss time of a mutation

In the same manner, the mean loss time of a mutation, i.e., the average number of generations for a mutation to be lost from the population, can be computed by

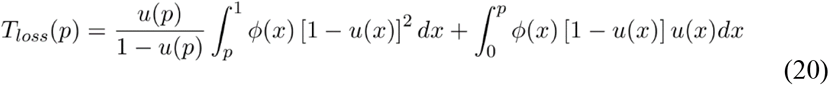

We discuss some special cases as follows.

Similarly, we have shown that the mean loss time of a selection-duality mutation given an initial frequency, *T*_*loss*_(*p*), is identical between the substitution neutrality and the FSI-neutrality. In the case of substitution neutrality, the mean loss time can be computed by the following formula,

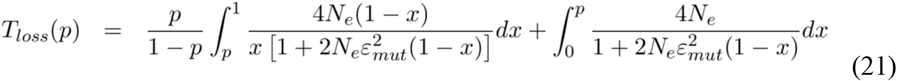

After some calculations, we show

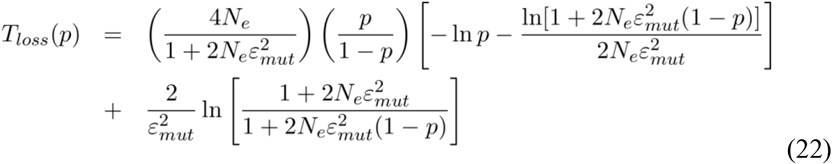

In the case of no FSI such that 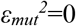, it is reduced to the classical result

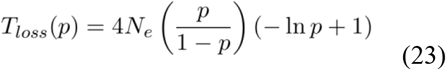

#### Mean loss time of a new mutation

In practice, we are more interested in the mean loss time of a new mutation with a very low initial frequency. Consider a new mutation with the initianl frequency 1/(2*N*), where *N* is the census population size. Under the substitution neutrality or the FSI-neutrality, we obtain

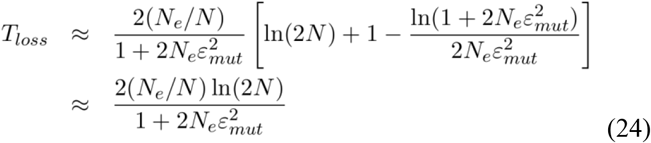

Hence, the mean loss time could be much short when FSI is strong.

While we show that mean loss time is identical between the substitution neutrality and the FSI neutrality, it is difficult to derive a formula for the mean loss time under the positive selection-duality. Nevertheless, in the case of new mutation when *p*=1/(2*N*) that is close 0, we approximately have

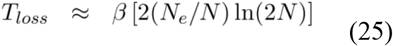

where β is chosen such that the approximate formula is identical to *T*_*loss*_ in the substitution neutrality or the FSI-neutrality, as given by

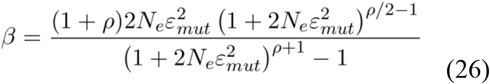

One may see the Appendix in details.

## Discussion

### The mean fixation generations of selection-duality mutations

The most striking finding of our study is that the mean fixation time of a mutant under FSI is identical between the substitution neutrality (*ρ* = −2) and the FSI-neutrality (*ρ* = 0). In the area between them, called the positive selection-duality, there is no analytical form for the mean fixation time, yet some approximate results have been derived. Numerical analysis showed that the difference is usually negligible. On the other hand, we have derived the mean fixation time at the generic neutrality (ρ = −4), which is slightly less than two other neutrality cases. The underlying mechanism that may cause those slight differences remains unclear. Putting together, one may conclude that FSI can speed up the fixation process of selection-duality mutations considerably, with roughly the same magnitude.

### The total time for de novo mutations to arise and fix in a finite population

The scenario of molecular evolution can be characterized by two evolutionary time scales. In addition to the number of generations taken for a mutant to fix in the population, the pace of molecular evolution also depends on how long it takes for such an allele to arise. We follow Glémin and Ronfort (2013) and define the total expected number *T*_*new*_ of generations for a mutation with frequency *p* to be fixed ultimately in the population, that is,

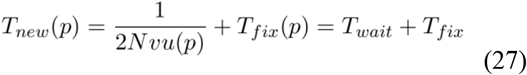

The first term on the right hand of Eq.(27) is the waiting time, *T*_*wait*_, the expected number of generations for the emergence of a mutation that will be fixed ultimately, where *v* is the mutation rate, *u*(*p*) is the fixation probability, and *N* is the census population size. The second term is the mean fixation time as defined above. For *de novo* mutation with the initial frequency *p*=1/(2*N*), the waiting time is the inverse of the substitution rate (λ). Without loss of generality, substitution rate can be symbolically written as *λ* = *vf* and the mean fixation time by *T*_*fix*_ = 4*N*_*e*_*g*, Eq.(27) can be written as follows

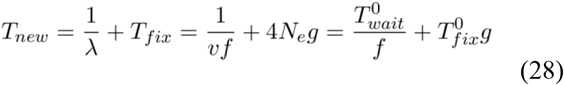

where 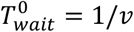 and 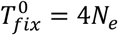 are the waiting time and the fix time of a strictly neutral mutation, respectively. For beneficial mutations, we have *f* > 1 and *g* < 1, suggesting that positive selection could decrease both waiting time and fixation time considerably. By contrast, for deleterious mutations, we have *f* < 1 and *g* > 1, suggesting that purifying selection could increase both waiting time and fixation time considerably. Intriguingly, selection-duality mutations under FSI reveals a mixed pattern: our previous analysis showed have *f* < 1 and *g* < 1, suggesting a prolong waiting time and a shortened fixation time. It appears that the mean fixation time *T*_*fix*_ of selectionduality mutations could is much less than the waiting time *T*_*wait*_ when FSI is nontrivial. Consequently, the nucleotide substitution between species can be unambiguously separated from the stage of population genetics, which can be modeled by a Markovchain process. How this finding would impact on our understanding of molecular evolution requires a further study.

### Selective sweeps and selection duality

In population genetics, a selective sweep refers to a process where a new, beneficial mutation rapidly increases in frequency and becomes fixed in a population driven by positive selection (Maynard-Smith and Haigh 1974). The ‘hitchhiking’ effects of selective sweeps could result in a considerable reduction of genetic variation in the genomic region surrounding the beneficial allele. The controversy is whether a reduced genetic diversity in a genome region must be a signature of ancient adaptive selections that had occurred in the evolutionary history of the genome.

The nature of rapid fixation of a selection-duality mutation when FSI is strong 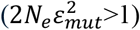 provides a new interpretation for the reduced genetic diversity region, as referred to selection-duality sweeps. While a detailed theoretical analysis is beyond the scope of this article, we present an intuitive approach as follows. It appears that the null hypothesis of selective sweeps is when a strictly neutral mutation is arising toward fixation by genetic drift. During the fixation time, the pre-standing genetic variation, (assumedly 4*N*_*e*_*v*) in the surrounding genomic region was swept, and meanwhile, new mutations were accumulating with a rate of *v* per generation. Since the mean fixation time (*T*_*fix*_) of a neutral mutation is 4*N*_*e*_ generations, the expected amount of accumulated mutation is 4*N*_*e*_*v*. Thus, fixation of a neutral mutation will not lead to a reduction of genetic variation in the surrounding region. This approach was first suggested by Gillespie (2000; 2001).

To explain an observed reduction of genetic variation, people invoked beneficial mutations whose fixation processes are much faster than neutral ones, without leaving sufficient time for accumulation of new mutations in the surrounding region. It is surprising that selection-duality mutations may also have experienced a rapid fixation process when FSI is strong. In the case of either fixation neutrality or FSI-neutrality, one can show that the ratio of *T*_*fix*_ to 4*N*_*e*_ is about 69.3%, 35.8% and 24.0% for 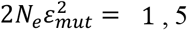, 5 and 10, respectively. Very roughly, our intuitive argument suggestion that the reduced genetic variation in the genome region surrounding a fixed selection-duality mutation is roughly given by

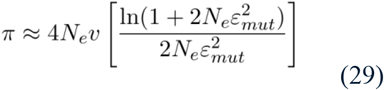

(here no recombination is assumed). Eq.(29) predicts that the reduction of genetic diversity at linked sites could be considerable in the case when 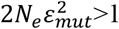, as shown by Fig.1. Our argument is, of course, intuitive, and more rigorous analysis will be the focus of future study.

**Fig. 1.**
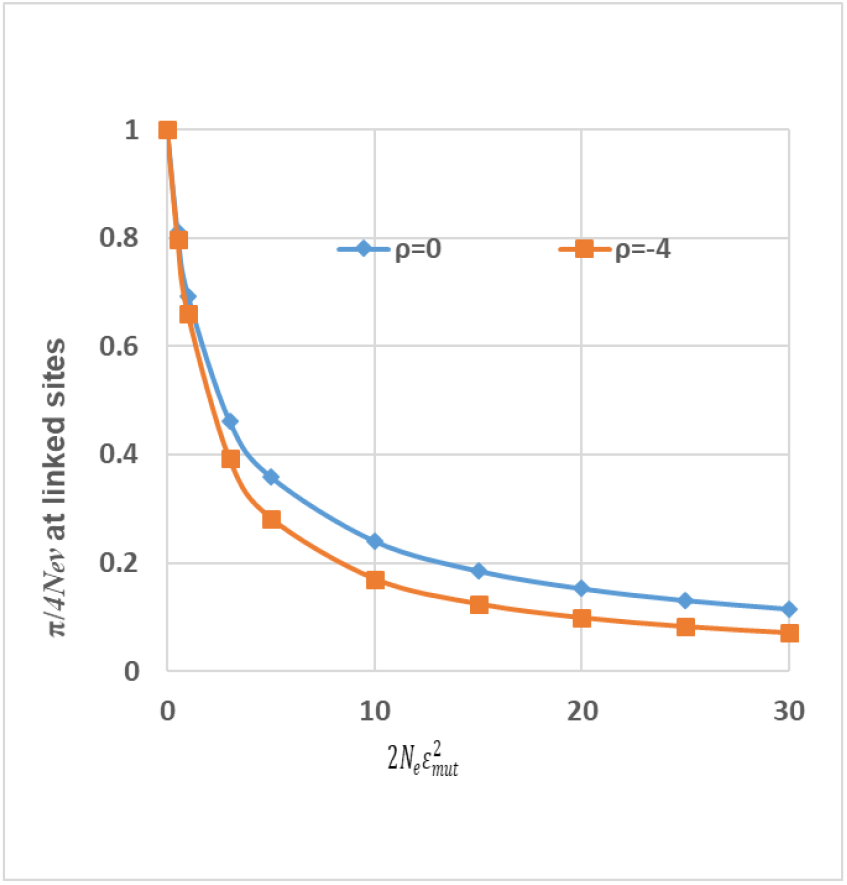
The expected genetic reduction at linked sites due to the fixation of a selection-duality mutation. The area of selection-duality is between the two lines: the substitution neutrality (*ρ* = 0) and the generic neutrality (*ρ* = −4).

## Methods

### Derivation of the mean fixation time

The mean fixation time, denoted by *T*_*fix*_(*p*), is the conditional expectation of the generations required for a mutant with the initial frequency *p* to be ultimately fixed in the population. Let *u*(*p,t*) be the probability that the mutant becomes fixed in the population by generation *t*, given the initial gene frequency *p*. Since the probability that the mutant gene is fixed at generation *t* is ∂*u*(*p, t*)/ ∂*t*, the average number of generations at which the gene is fixed is given by

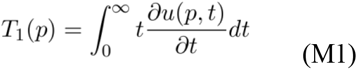

As shown by Crow and Kimura (1970); also in the Appendix, we obtain the following differential equation

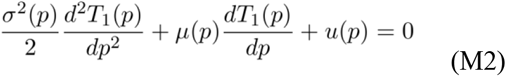

with the boundary conditions *T*_1_(0) = 0 and *T*_1_(1) = 0. Note that *T*_1_(*p*) includes the case when a mutant actually got eliminated, which should be excluded. If the eventual probability of fixation of the mutant gene is *u*(*p*), the mean fixation time *T*_*fix*_(*p*) is then defined by

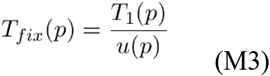

One may solve Eq.(M2) under *T*_1_(0) = 0 and *T*_1_(1) = 0, and obtain Eq.(7) after applying Eq.(M3).

### Calculation of the mean fixation time under selection-duality

Plugging Eq.(2) upon Eq.(8), we obtain the general expressions of *ϕ*(*x*) and *u*(*x*), as given by

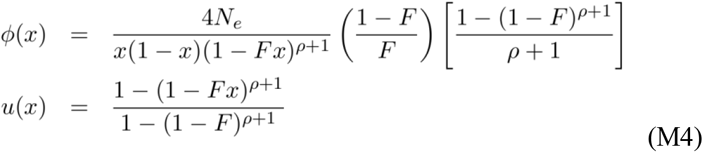

respectively. Given a special value of ρ=0, −2 or −4, it is straightforward to calculate *ϕ*(*x*) and *u*(*x*) and then apply to Eq.(7) or Eq.(8) to obtain *T*_*fix*_(*p*) or *T*_*fix*_.

## Appendix

### A general theory of mean fixation time

Let *u*(*p,t*) be the probability that the mutant gene frequency becomes fixed in the population by generation *t*, given that the initial gene frequency is *p*. Since the probability that the mutant gene is fixed at generation t is ∂*u*(*p, t*)/ ∂*t*, the average number of generations at which the gene is fixed is given by

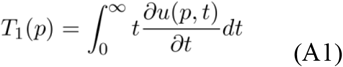

Note that the Kolmogorov backward equation of *u*(*p, t*) is given by

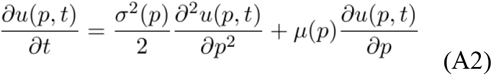

Differentiating each term of Eq.(A2) with respect to *t*, we have

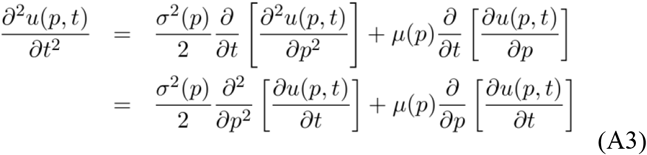

Next, multiplying both sides of Eq.(A3) by *t*, and integrating with respect to *t* from 0 to ∞, we have

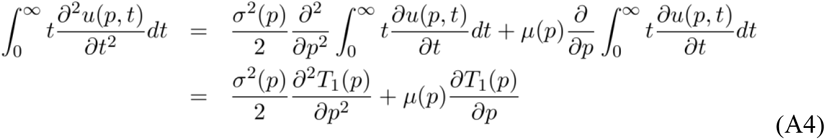

Meanwhile, the left hand of Eq.(A4) can be further simplified by

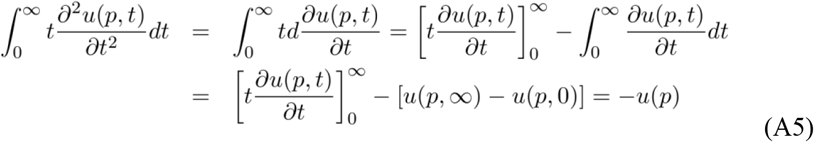

where we have assumed that is *t* ∂*u*(*p, t*)/ ∂*t* vanishes at *t* → ∞, *u*(*p*, 0) = 0, and *u*(*p*) = *u*(*p*, ∞). From Eq.(A4) and Eq.(A5), we therefore have the following differential equation

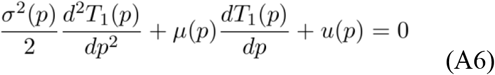

with the boundary conditions *T*_1_(0) = 0 and *T*_1_(1) = 0. Though the solution of Eq.(A6) is solvable, *T*_1_(*p*) includes the event in which the mutant gene is lost from the population. Hence, if the eventual probability of fixation of the mutant gene is *u*(*p*), the mean fixation time is then defined by

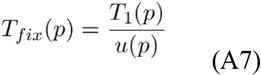

Together, we obtain

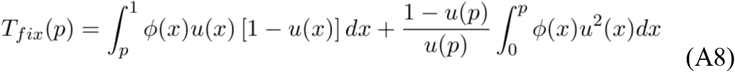

where *ϕ*(*x*) and *u*(*x*) are given by

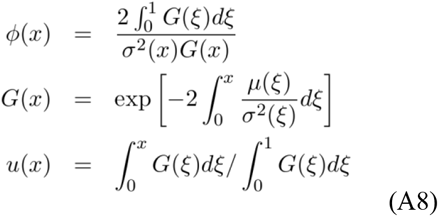

respectively.

The average number of generations for a mutant gene to be lost from the population can be obtained in the same way, where *T*_0_(*p*) is defined by

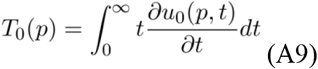

After removing the event of being ultimately fixed, the mean loss time is then defined by

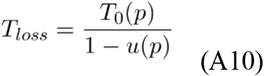

and can be calculated as follows

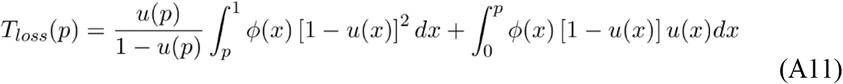

#### Equivalence between fixation neutrality and FSI-neutrality

An intriguing result is that both *T*_*fix*_(*p*) and *T*_*loss*_(*p*) are identical between the fixation neutrality (ρ = 0) and the FSI-neutrality (ρ = −2). A concise proof is given below. Based on Eq.(A8), one can easily verify that the following three formulas

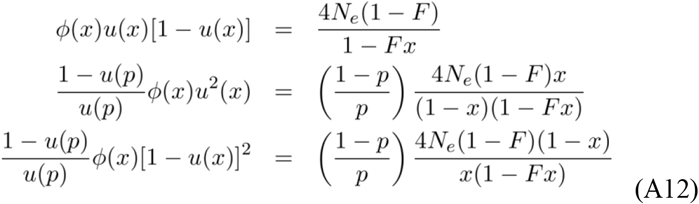

hold in both cases ρ = 0 and ρ = −2. We thus proof our claim according to Eq.(A8) and Eq.(A11).

#### Mean fixation (or loss) time of a new mutation

We first consider the mean fixation of a new mutation by letting the initial frequency to be aero, that is, 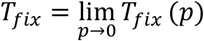. When *p* → 0, it can be shown that *u*(*p*)=0, *u*^**′**^(*p*) = 1 and *ϕ*(*x*) < ∞. It follows that the limit of the second integral on the right hand of Eq.(M8) can be calculated by using the L’Hopital’s rule, i.e.,

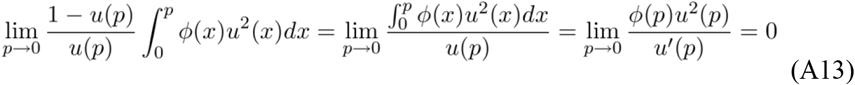

Therefore, the mean fixation time of a new mutation is given by

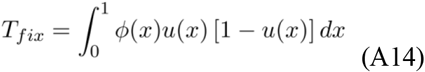

Next we consider the mean loss time of a new mutant by letting the initial frequency to be 1/(2*N*), where *N* is the census population size. i.e., 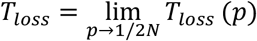. It appears that the second term on the right hand of Eq.(A11) approaches to 0 as *p* → 0.

Thus, *T*_*loss*_ can be approximated calculated by

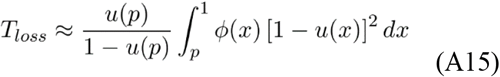

#### Approximate formulas of *T*_*fix*_ when −2 ≤ *ρ* ≤ 0

First we rewrite *T*_*fix*_ in the form as follows

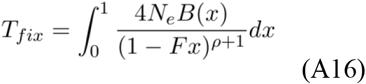

where *B*(*x*) is given by

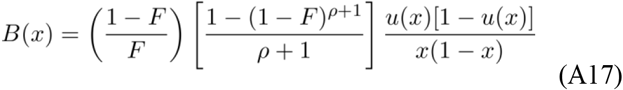

Mathematically, one can verify that *B*(*x*) is bounded in the range (0, 1). After some numerical analysis, we found that in the range of −2 ≤ *ρ* ≤ 0, *B*(*x*) can be roughly approximated by

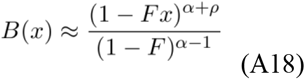

where α is a positive constant that remains to determine. Putting together, we obtain

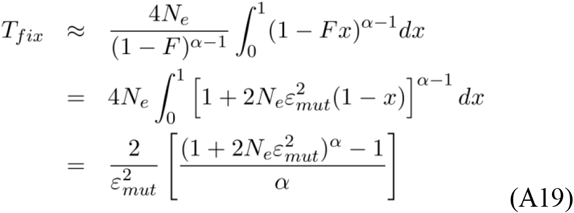

Finally, we determine constant α by the condition that at the two boundaries, *ρ*=0 or *ρ*=−2, the approximate formula of *T*_*fix*_ must be identical to the accurate one. One may then claim that α=0 when *ρ*=0 or *ρ*=−2. Tentatively, we choose

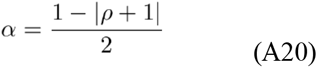

that satisfies the condition that α=0 when *ρ*=0 or *ρ*=−2. It appears that the maximum value of α is ½ when *ρ*=−1, in this case we have

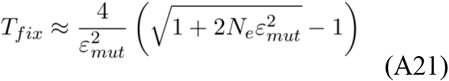

We therefore conclude that, under the positive selection-duality, the mean fixation time is, approximately, in the following range

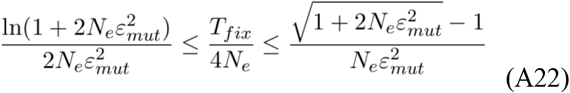

#### Approximate formulas of *T*_*fix*_ when −2 ≤ *ρ* ≤ 0

It has been shown (see later) that the mean fixation time of a mutation with *ρ*=−3 is given by

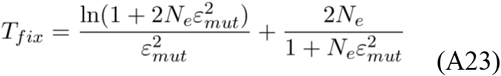

When 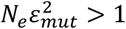, Eq.(A23) can be approximated by

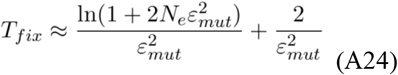

After comparing formulas of *T*_*fix*_ between Eq.(12) for *ρ*=−2, Eq.(23) for *ρ*=−3 and Eq.(14) for *ρ*=−4, we thus propose an approximate formula for −4< *ρ*<−2, that is,

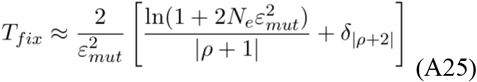

where *δ*_|*ρ*+2|_ is an index function: *δ*_|*ρ*+2|_ =0 when |ρ + 2| = 0, otherwise *δ*_|*ρ*+2|_ =1.

#### Approximation of *T*_*loss*_

While we show that the mean loss time is identical between fixation neutrality and FSI neutrality, it is difficult to derive a formula for the mean loss time under the positive selection-duality. Nevertheless, in the case of new mutation when *p*=1/(2*N*) that is close 0, we approximately have

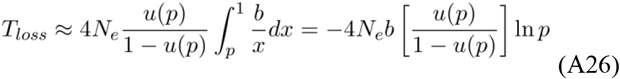

Where *b* is a constant to determine. It is known that as *p* is close to 0, we have

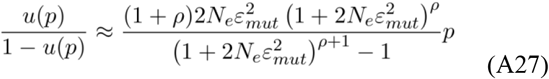

Together with *p*=1/(2*N*), we obtain

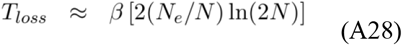

where

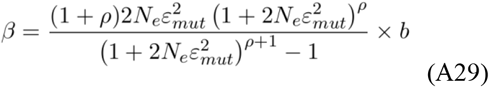

In particular, we choose a specific *b* as

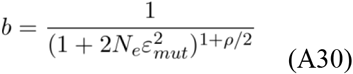

Such as the approximate formula is identical to *T*_*fix*_ in the fixation neutrality or FSI-neutrality. Therefore,

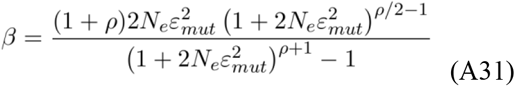

#### Two examples for the mean fixation time calculation

##### *Negative selection-duality* (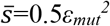 *or ρ*=−3)

First of all, we calculate the those quantities as follows

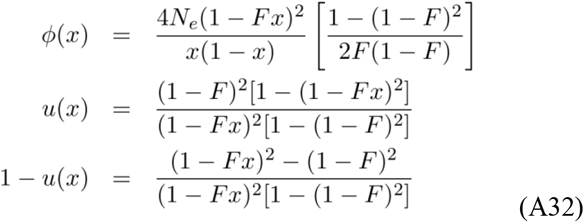

It follows that

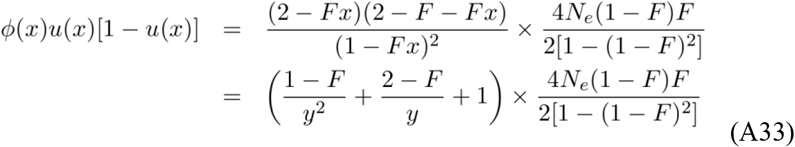

where *y*=1-*Fx*. We then derive *T*_*fix*_ by

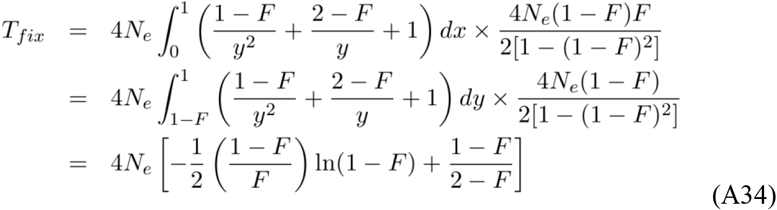

##### *Genetic neutrality* 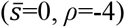

The calculation of *T*_*fix*_ in the case of generic neutrality is similar to the case shown above, in spite of more algebras. First we calculate

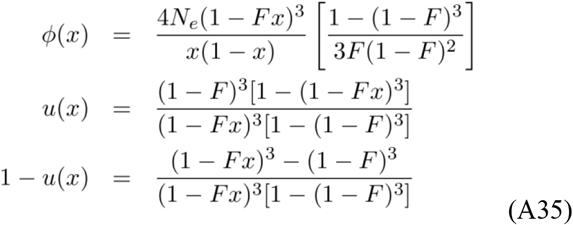

Let *y*=1-*Fx*, and constant *B* by

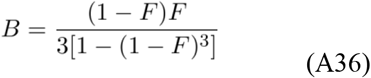

One may carry out the calculation of product *ϕ*(*x*)*u*(*x*)**[1** − *u*(*x*)] and simplification as follows

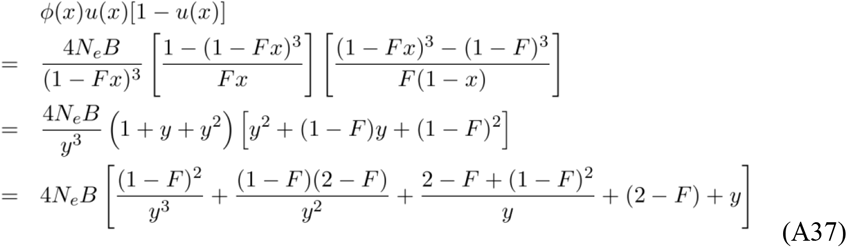

And finally, we have

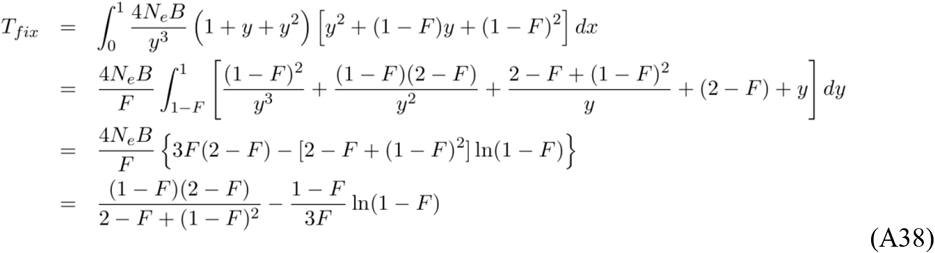

